# An R package for divergence analysis of omics data

**DOI:** 10.1101/720391

**Authors:** Wikum Dinalankara, Qian Ke, Donald Geman, Luigi Marchionni

## Abstract

Given the ever-increasing amount of high-dimensional and complex omics data becoming available, it is increasingly important to discover simple but effective methods of analysis. Divergence analysis transforms each entry of a high-dimensional omics profile into a digitized (binary or ternary) code based on the deviation of the entry from a given baseline population. This is a novel framework that is significantly different from existing omics data analysis methods: it allows digitization of continuous omics data at the univariate or multivariate level, facilitates sample level analysis, and is applicable on many different omics platforms. The divergence package, available on the R platform through the Bioconductor repository collection, provides easy-to-use functions for carrying out this transformation. Here we demonstrate how to use the package with data from the Cancer Genome Atlas.

## Introduction

The technologies that provide us with high-dimensional omics data continue to advance at a rapid rate. Particularly in the last decade, the available modalities of omics data have considerably expanded and now include, among others, coding and noncoding RNA expression, micro RNA expression, protein expression, epigenetic profiling related to histones and CpG methylation, copy-number and mutation profiling. As a result, the analysis of multi-modal omics data has become indispensable in many domains of biological and medical research.

Whereas the quantity and quality of available data has appreciably increased, the results of research based on such data are often not reliable, robust and replicable. A key challenge has been to quantify the level of variability and diversity in omics profiles in a given population, and to separate normal and technical variability from abnormal variability indicative of a biological property such as disease.

Recently we have introduced divergence analysis as a method for simplifying high-dimensional omics data for bioinformatic analyses [1]. This method conceptually parallels the widespread use of deviation from normality as a disease marker in clinical testing, such as a blood based prostate specific antigen (PSA) test [2,3]. Given a high-dimensional omics data profile, it can be converted to a binary or ternary string of the same length, where each value now indicates how the original value diverges from a baseline or reference population. The features of interest may be univariate such as a gene, a CpG site, or a protein, in which case the original profile is converted to one of three labels 0, −1 and 1 indicating whether the level of a feature is, respectively, within the reference range for that feature, or below or above that range. Or, the features of interest may be multivariate, such as sets of genes representing pathways of interest. In this case, divergence coding is binary, where a 0 or 1 for a multivariate feature indicates, respectively, that the set of feature is inside or outside the support of the multivariate, baseline distribution.

We have prepared the ‘divergence’ R package [4] offered through the bioconductor package repository [5] (https://www.bioconductor.org/packages/release/bioc/html/divergence.html) which provides functionality for divergence based computations.

Divergence coding can be used for a wide variety of types of data analysis: class comparison and prediction, feature selection, regression, combination of omics modalities, and many others. Here we present how to use this package and showcase the many different ways in which the divergence coding can be manipulated to perform interesting analyses. In the following we use breast normal and tumor samples from the TCGA project [6] for which RNA-seq expression profiles as well as methylation profiles are provided. First we use some basic examples to illustrate the workflow, followed by more complex types of data analyses that combine multiple omics modalities, and demonstrate how to use divergence in both the univariate and multivariate settings. A subset of this data is available with the R package.

## Methods

We may summarize the divergence method as follows. Before computing the baseline range (univariate) or support (multivariate) and the resulting divergence coding, a rank-transformation is applied to all data. Consider an omics sample represented by a vector *X* = *(X_j_),j* = 1..*m, m* being the dimensionality of the omics profiles. The following transformation is applied which converts the data to a normalized rank, with the minimum being zero.

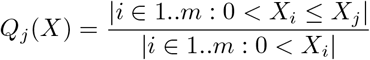

Then *Q_j_* (*X*) ∈ [0,1]. The zero minimum is particularly useful in preserving the zero valued mass usually observed in many omics data modalities, such as RNA-Seq.

Now consider a multivariate feature indexed by *S* - i.e. a subset of the given m features (the univariate scenario is a special case when |*S*| = 1). We will denote the corresponding subset of a given omics profile X following the quantile transformation as Q(X)^S^. Suppose we have *n* such profiles that constitute the baseline group. We estimate the baseline support as follows: given a parameter *γ ∈* [0,1], we compute *l* which is the floor of *nγ*. Then from each baseline sample *k*, if *r_k_* is the distance from *Q(X)^S^* to it’s *l^th^* nearest neighbor in the multivariate feature space. If we denote the sphere around *Q(X)^S^* of radius *r_k_* as 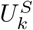 then the support of the baseline is the union of the regions covered by these spheres around each baseline sample, which we denote as *Û^S^*

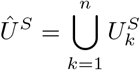

Following the baseline estimation in this manner, for any given omics profile *X*, the divergence coding *Z*(*X*)^*S*^ can be computed as:

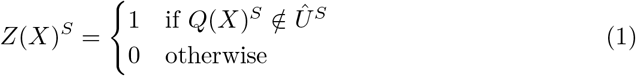

For the univariate scenario, the support is the union of a series of intervals. However, we apply a further simplification by replacing these with a single interval spanning the lowest end to the highest end of these intervals. Accordingly the divergence coding becomes ternary: *Z(X)^j^* ∈ {−1, 0,1} with −1 indicating a value below the baseline interval for feature *j*, 1 indicating a value above the baseline interval, and 0 indicating no divergence.

In practice we use two more parameters, *α* and *β*. The *β* parameter is used for allowing a certain number of outliers to be exempted from the baseline. Given the radii of the spheres rι..r_n_, let 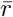 be the (1 − *β)^th^* percentile of these values. Then we select only the spheres with 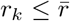 to compose the baseline. Once the baseline is computed, we can compute the divergence coding for the baseline samples; then *α* is the average proportion of divergent features (multivariate or univariate) among the baseline cohort. As discussed in the following section, we specify *α* and *β* and then select the *γ* value that fits these specified parameters. However the functionality for estimating a baseline for a specified γ parameter is available in the package as well.

For a more detailed description of the method, see [1].

## Results

### Univariate Workflow

To carry out an analysis based on divergence, the first step is to determine the case and control samples. The control cohort here will be used for computing the baseline interval for each feature, and we will refer to it as the baseline cohort or baseline group. Once the baseline is computed, the divergence values for the case cohort will be computed with reference to the baseline.

What data should be used as the baseline cohort depends on the problem at hand and needs careful investigation based on the biological or clinical questions of interest that are being investigated. In many scenarios of disease based data, and in particular cancer, normal samples would be a good choice. The more normal samples available, the more robust the baseline will be.

All samples, both in the case cohort and the baseline cohort should have the same features - for example genes or microarray probes, and be from the same platform (e.g. RNASeq or a specific microarray platform). Ideally the baseline cohort should be from the same experiment to avoid study-specific batch effects as much as possible. In general we suggest at least 20 samples be available to compute the baseline.

The divergence algorithm applies a scaled rank transformation to all samples before the digitization of the data. The package provides functionality for the user to compute this rank transformation manually, or it can be set to compute internally when the divergence computation is requested by the user. No other normalization procedures are applied, and the user may apply any normalization procedures beforehand as necessary.

There are three parameters involved in the divergence computation: *α, β* and *γ*. All parameters are in the (0, 1) range. The *β* parameter adjusts the support to account for a certain percentage of outliers included in the baseline data, and the *γ* parameter provides a way to widen or tighten the support around each baseline sample. The closer *γ* is to 1, the further the support around each normal sample will extend. For a given support, the *α* value is simply the average number of divergent features per sample for the baseline cohort.

In usage, we usually provide the *α* and *β* values and a range of possible *γ* values and let the package find the most appropriate *γ* value out of these for the given a and *β* values. By default, the package uses *α* = 0.01, *β* = 0.95, and *γ* ∈ {0.01, 0.02,…, 0.09,0.1, 0.2,…, 0.9} as a list of candidate *γ* values. Thus if you use these default values, the package will consider 95% of the baseline cohort to be included in the support and find the smallest possible *γ* value from the given list that will provide the average number of divergent features per sample in the baseline cohort to be 1% or less. For more details, see [1].

We use RNA-Seq data spanning 20530 genes from the TCGA Breast Cancer dataset [6,7] for the following analysis. This data consists of 1097 tumor samples and 114 normal samples, the latter which we will use as our baseline cohort.

Using the default parameters available, we compute the digitized ternary divergence format for the tumor samples with respect to the normal baseline of 114 samples. The algorithm selects a *γ* value of 0.3, which yields an *α* value of 0.007 which is below the *α* threshold of 1% which was specified as the default.

The tumor data contains 601 ER+ and 179 ER− samples. We can see whether the number of divergent genes per sample are similar between the two ER groups or not (Figure 1).

**Fig 1.**
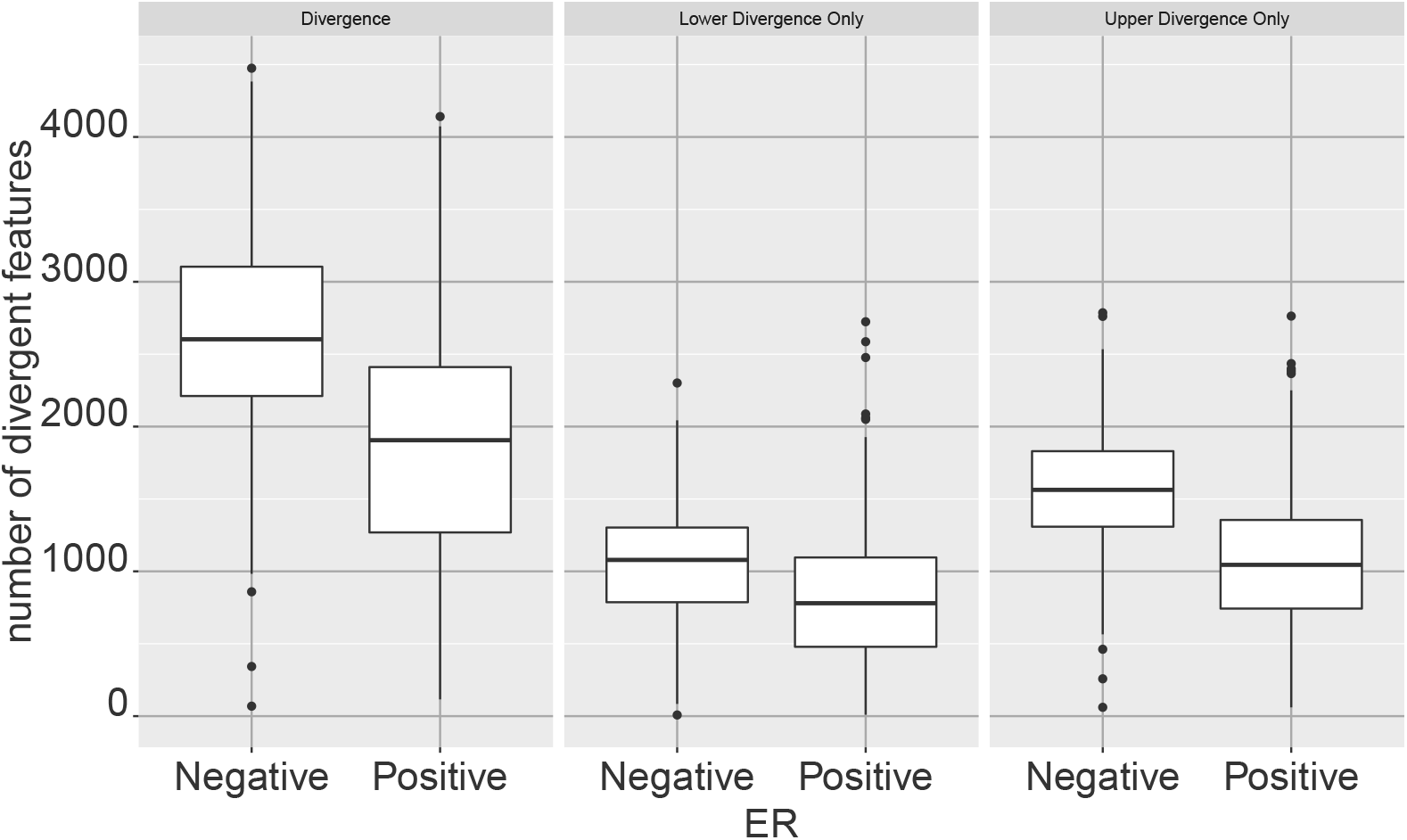
Number of divergent features as a measure of sample divergence. The size of the divergent set in each sample - i.e. the set of features that are divergent in a given sample - can be uses as a measurement of the divergence of a sample. This allows the comparison of sample level divergences between sample groups. When digitized with respect to a normal breast basenline, the boxplots compare the number of divergent features per sample between ER+ and ER− breast tumors. Given the higher risk associated with ER-breast tumors, we expect to observe higher divergence from normality compared to ER+. The same trend is observed if the consideration is limited to the number of upper or lower divergent features only.

As another example of using the size of the divergent feature set as an indicator of the divergence of the sample, we can compare it against clinically obtained covariates such as the percentage of normal and tumor cells found in the sample. We see that these percentages track with sample level divergence (Figure 2).

**Fig 2.**
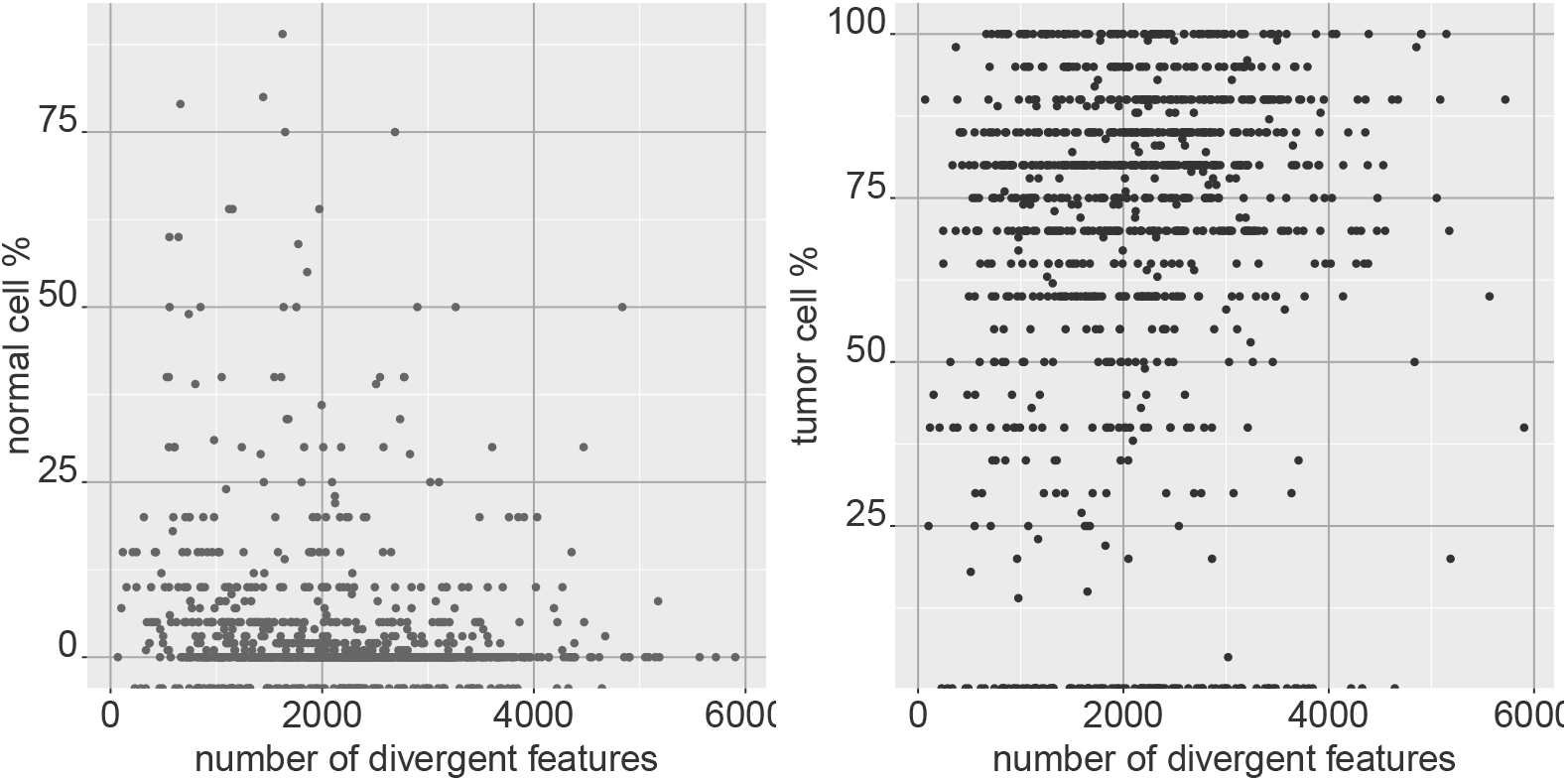
Comparing sample divergence with clinical covariates. Sample divergence of breast tumor samples (as measured by the size of the divergent set in each sample), with respect to a normal breast baseline, shows negative correlation (spearman correlation = −0.266) with the normal cell percentage estimate of the sample, and positive correlation (spearman correlation = 0.167) with the tumor cell percentage estimate of the sample, illustrating that sample divergence is indicative of the sample level deviation from normality.

Table 1 shows the correlations obtained with the percentages of normal, tumor, and stromal cells provided from pathologist observations in the TCGA data. A similar comparison can be made with the cell type enrichment scores provided by xCell, a method which provides a score for gene expression samples suggesting the level of enrichment in the sample for different cell types [8]. As see in table 1, the number of divergent genes in breast tumor samples show high correlation with the enrichment scores provided for many cell types. These results indicate that some of the divergence observed is reflective of the cell type heterogeneity in the specimen.

**Table 1.** Correlation between cell type signatures and divergence score. The divergence score (number of divergent genes in a sample) for breast tumor samples based on gene expression is correlated with the scores from the xCell tool reflecting enrichment in the sample for different cell types. The 20 cell types with the largest spearman correlations are shown. Also shown are correlations between the divergence score with the normal, tumor, and stromal cell percentages provided in the TCGA clinical covariate data.

| cell type | correlation |
| --- | --- |
| Type 1 helper (Th1) cells | 0.779 |
| Megakaryocyte-erythroid progenitor (MEP) | 0.636 |
| Plasma cells | 0.594 |
| Type 2 T helper (Th2) cells | 0.457 |
| Natural killer T cells (NKT) | 0.440 |
| Osteoblast | 0.417 |
| pro B-cells | 0.352 |
| Common lymphoid progenitor (CLP) | 0.342 |
| Hematopoietic stem cells (HSC) | -0.636 |
| Mesangial cells | -0.589 |
| Chondrocytes | -0.552 |
| Endothelial cells | -0.528 |
| Adipocytes | -0.505 |
| Hepatocytes | -0.501 |
| Megakaryocytes | -0.493 |
| Fibroblasts | -0.474 |
| Lymphatic endothelial cells | -0.472 |
| Conventional dendritic cells (cDC) | -0.430 |
| Mast cells | -0.410 |
| CD4+ naive T-cells | -0.354 |
| normal cell % | -0.266 |
| tumor cell % | 0.167 |
| stromal cell % | -0.170 |

To perform a differential expression analysis at the feature level for digitized divergence data, the package provides a *χ*^2^ test functionality, which we can use to perform a *χ*^2^ test for each gene between ER+ and ER− samples. Even simpler, we can merely compute the divergent probability of each gene over the ER+ and ER− samples respectively. Note that the divergent probability of a feature over a group of samples is simply the number of samples for which the feature is non-zero in the divergence space divided by the number of samples.

Figure 3 shows the divergent probability of each gene for ER+ and ER− samples. The samples in blue are genes that are significant under a Bonferroni-adjusted *P* ≤ 0.05 threshold from the *χ^2^* test. The ESR1 gene, which can be considered a marker for ER status, is highly significant and is colored in red. Table 2 shows the top ten genes by rank of *χ*^2^ test p-value.

**Fig 3.**
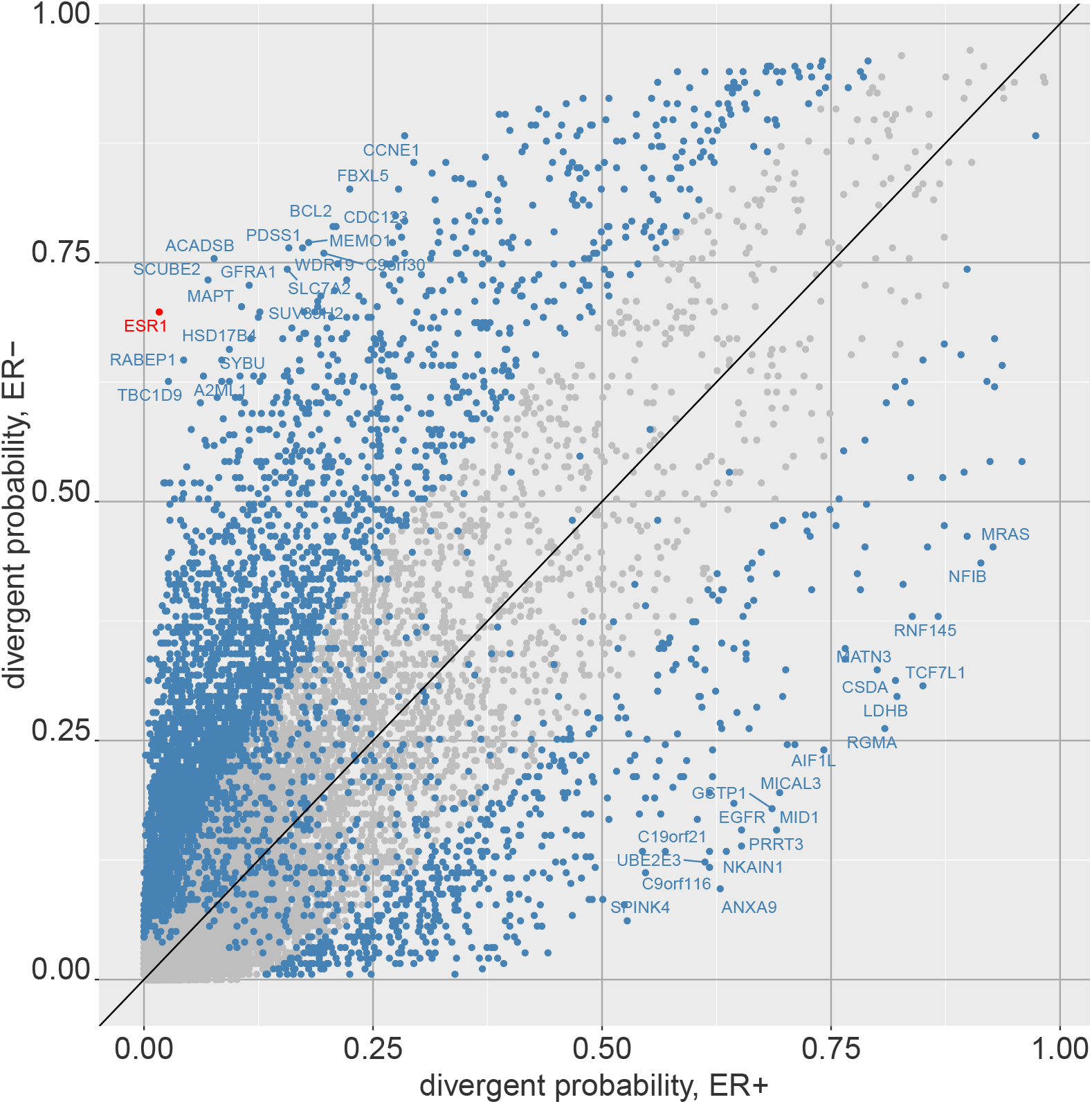
Differentially divergent features between two phenotypes. The probability of divergence for each feature is plotted for two sub-groups of breast tumors, ER+ and ER−. Genes far from the diagonal are likely to have highly different divergence patterns between the two groups (e.g. ESR1). Alternatively, a *χ*^2^ test can be performed for each feature between the two groups, and points in blue indicate genes that are *χ*^2^ test p-value significiant at a bonferroni adjusted *P* ≤ 0.05 threshold.

**Table 2.** 10 Most differentially divergent genes between ER+ and ER− breast tumor samples.

| gene | divergent<br>probability<br>ER+ | divergent<br>probability<br>ER- | Chi-squared<br>test statistic | p-value |
| --- | --- | --- | --- | --- |
| ESR1 | 0.0166 | 0.6983 | 443.0622 | $P < 10^{-6}$ |
| TBC1D9 | 0.0266 | 0.6257 | 356.4973 | $P < 10^{-6}$ |
| ACADSB | 0.0765 | 0.7542 | 351.6167 | $P < 10^{-6}$ |
| SCUBE2 | 0.0699 | 0.7318 | 346.3227 | $P < 10^{-6}$ |
| RABEP1 | 0.0433 | 0.648 | 334.7269 | $P < 10^{-6}$ |
| PSAT1 | 0.0616 | 0.6034 | 314.8622 | $P < 10^{-6}$ |
| CXorf61 | 0.0216 | 0.5196 | 286.9675 | $P < 10^{-6}$ |
| EN1 | 0.0233 | 0.5196 | 291.2844 | $P < 10^{-6}$ |
| A2ML1 | 0.0649 | 0.6313 | 278.4052 | $P < 10^{-6}$ |
| SKP1 | 0.0516 | 0.5754 | 279.7753 | $P < 10^{-6}$ |

We can observe how these genes track with the ER level by comparing the expression level to that of ESR1 expression, both in the regular expression value space and in the divergence space. Figures 4 and 5 show the values of ESR1 expression among ER+ and ER− breast tumor samples against two of the genes highly differentially expressed between the two sample groups, ACADSB and PSAT1, in the regular expression space (log_2_ transcripts per million) and in the divergence space.

**Fig 4.**
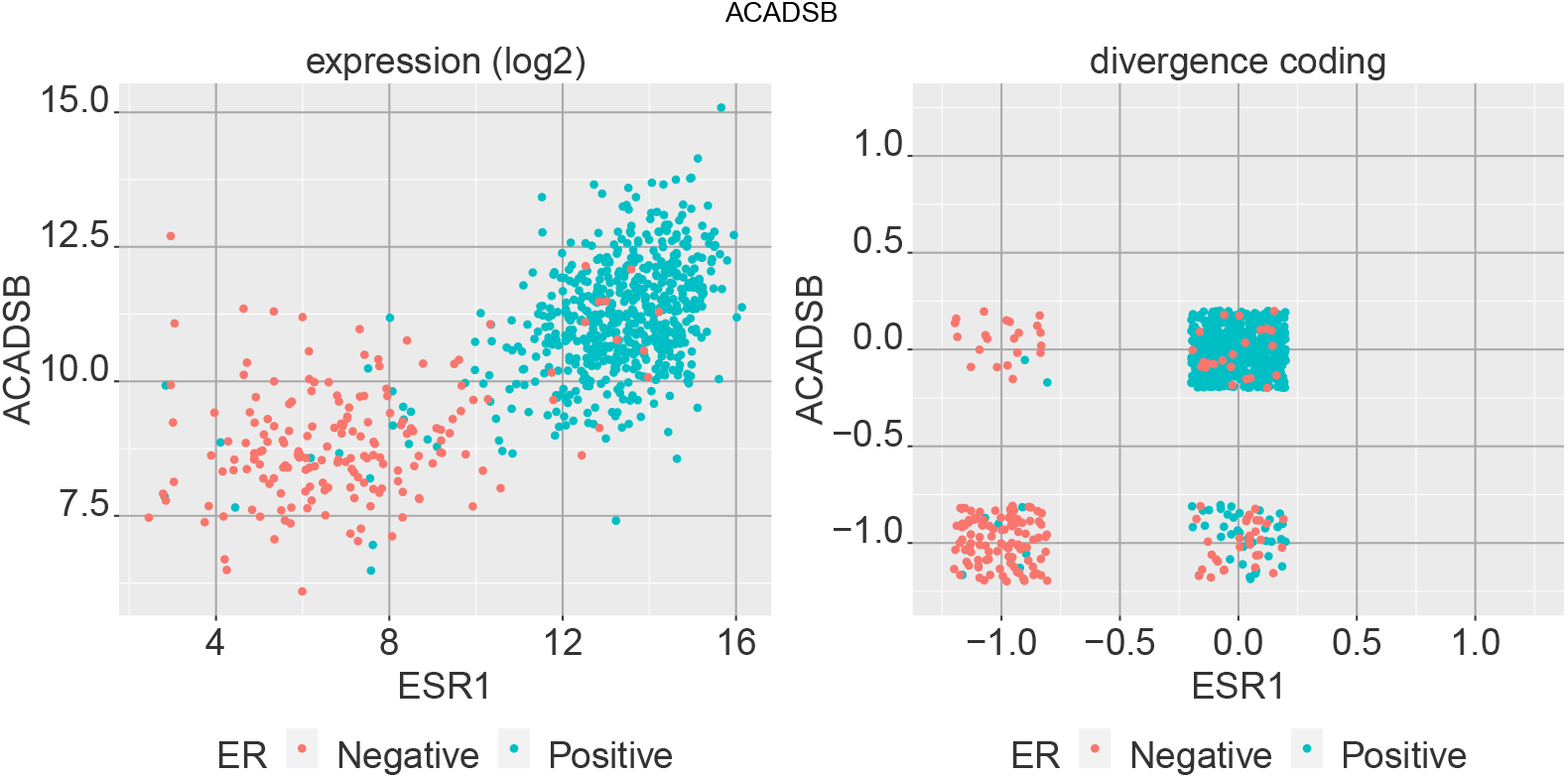
Comparing two genes in expression space and divergence space. Comparison of ACADSB and ESR1 gene expression for ER+ and ER− breast tumor samples in regular expression space (log_2_ TPM) and the digitized divergence space. ESR1 is an indicator of ER status and ACADSB shows to be highly differentiated between the two groups as well.

**Fig 5.**
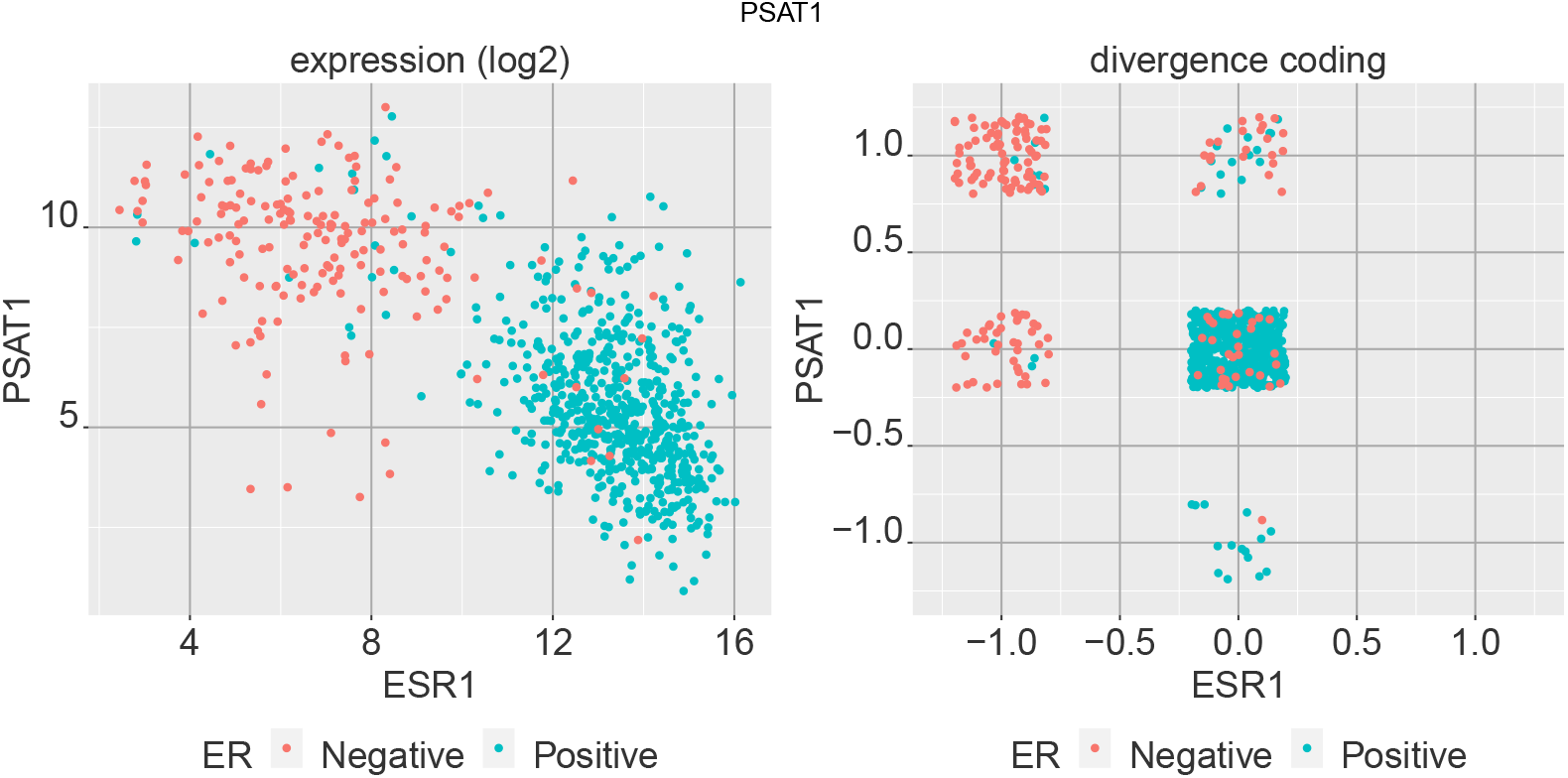
Comparing two genes in expression space and divergence space. Comparison of PSAT1 and ESR1 gene expression for ER+ and ER− breast tumor samples in regular expression space (log_2_ TPM) and the digitized divergence space.

### Multivariate Workflow

In the multivariate scenario, the features of interest are composite - that is, they are sets of univariate features, for example a gene set indicating a disease specific signature or a pathway. The workflow is similar to that of the univariate case, with the exception that the divergence coding will be either 0 or 1 - that is, it indicates whether the feature is divergent or not, whereas for the univariate case it provided a direction of divergence as well.

With the same breast gene expression data from TCGA which we have used so far, in the following we examine the divergence coding obtained for the KEGG pathways from MSIGDB gene set collection [9]. Computing the divergence coding is similar to the univariate case, except that the multivariate features need to be specified. As with the univariate case, the reference sample data and the samples for which the divergence coding need to be computed are provided in matrix form, along with a range of *γ* values to choose from, a *β* value, and the required *α* threshold. The multivariate features are provided as a list, where each element of the list is a vector of univariate features which are available in the data matrices provided. In the example we present, the same TCGA sourced RNA-Seq expression data used previously are used. The KEGG pathway list has 186 gene sets, where each set ranges from 10 to 389 gene symbols. These gene symbols are available in the RNA-Seq expression data and will be used to estimate the baseline support from the matched normal samples and the placement of the tumor samples either within or outside the support in the high dimensional space representing each KEGG gene set. The output will be a matrix comprising of the binary divergence coding for each KEGG pathway and each tumor sample.

Given that the divergence coding is a highly simplified representation, we can simply observe it in terms of either the features or the samples and observe the proportion of values in each divergence state. Figure 6 shows the sample proportions for some of the KEGG pathways, and similarly at the sample level in figure 7.

**Fig 6.**
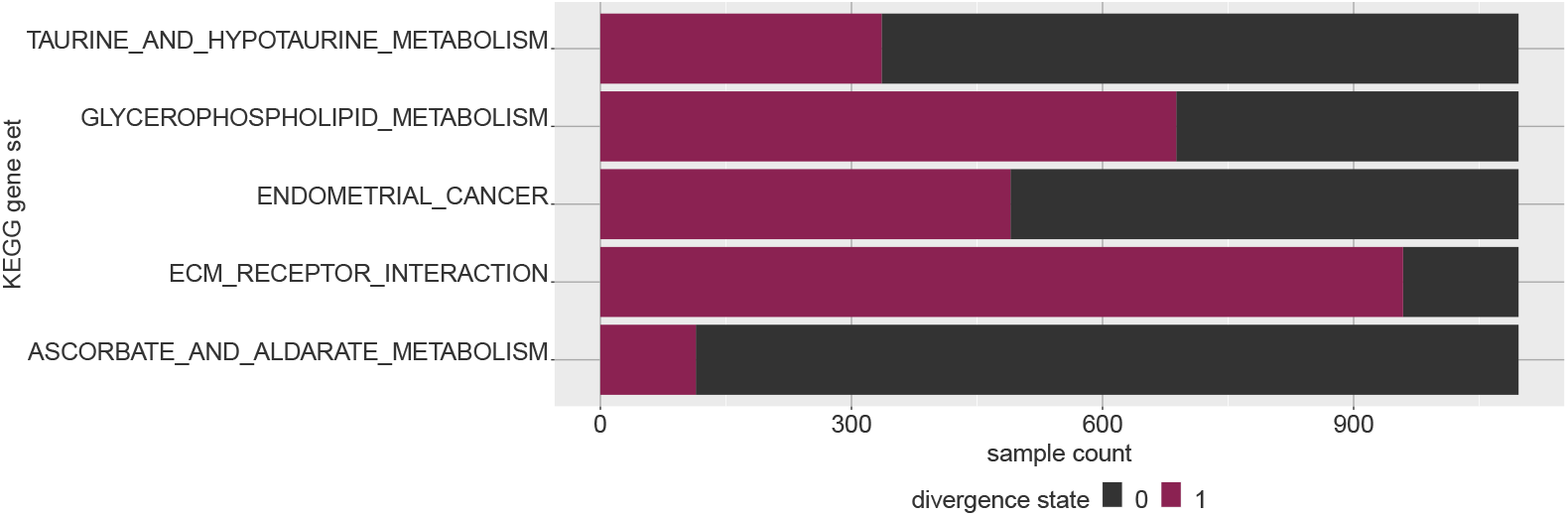
Visualizing multivariate divergence by feature. Given that multivariate divergence results in a binary digitization, for each feature we can observe how many samples are in a given state. Multivariate features (which are sets of genes, in this case) NOTCH signaling pathway and GNRH signaling pathway, for example, are much more likely to be divergent compared to the other features shown.

**Fig 7.**
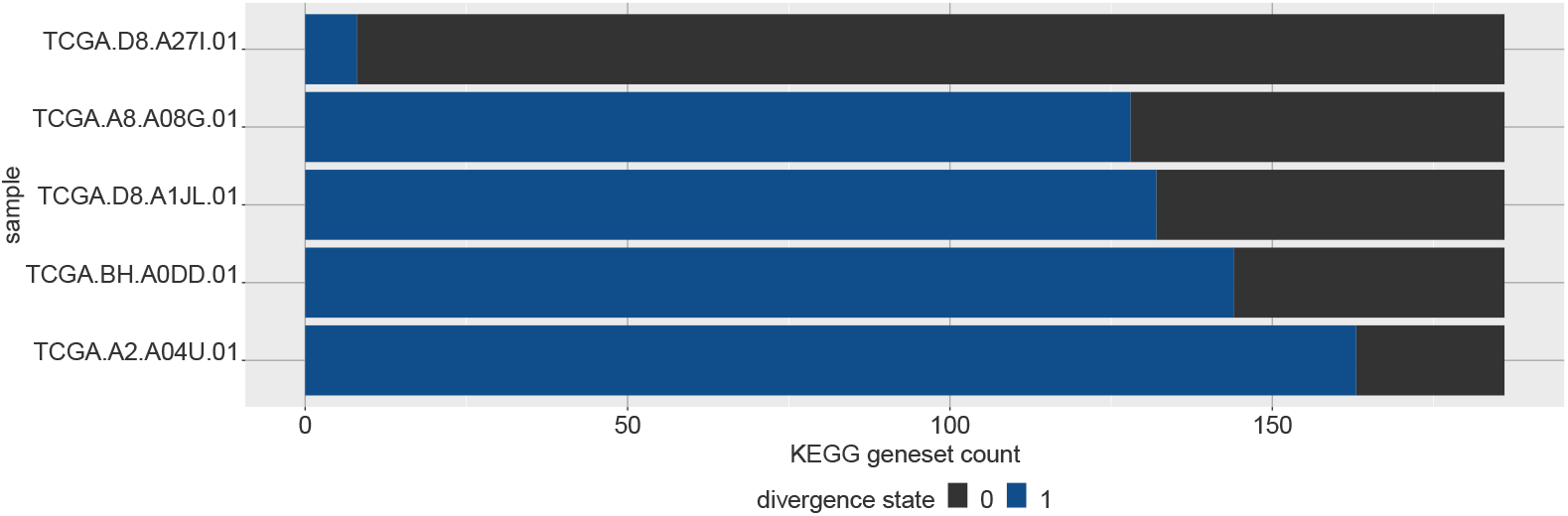
Visualizing multivariate divergence by sample. For a given set of multivariate features, we can see how likely they are to be divergent in a given sample. Here five selected breast tumor samples are ordered by how many gene sets in the KEGG gene set collection are divergent in each sample, with respect to a normal breast baseline.

As before, we can apply a *χ*^2^ test to each pathway between the ER+ and ER− sample groups to identify which pathways are highly differentially divergent between the two groups (Figure 8). The pathways that meet a *P* ≤ 0.05 cutoff after Bonferroni adjustment for multiple testing are shown in blue.

**Fig 8.**
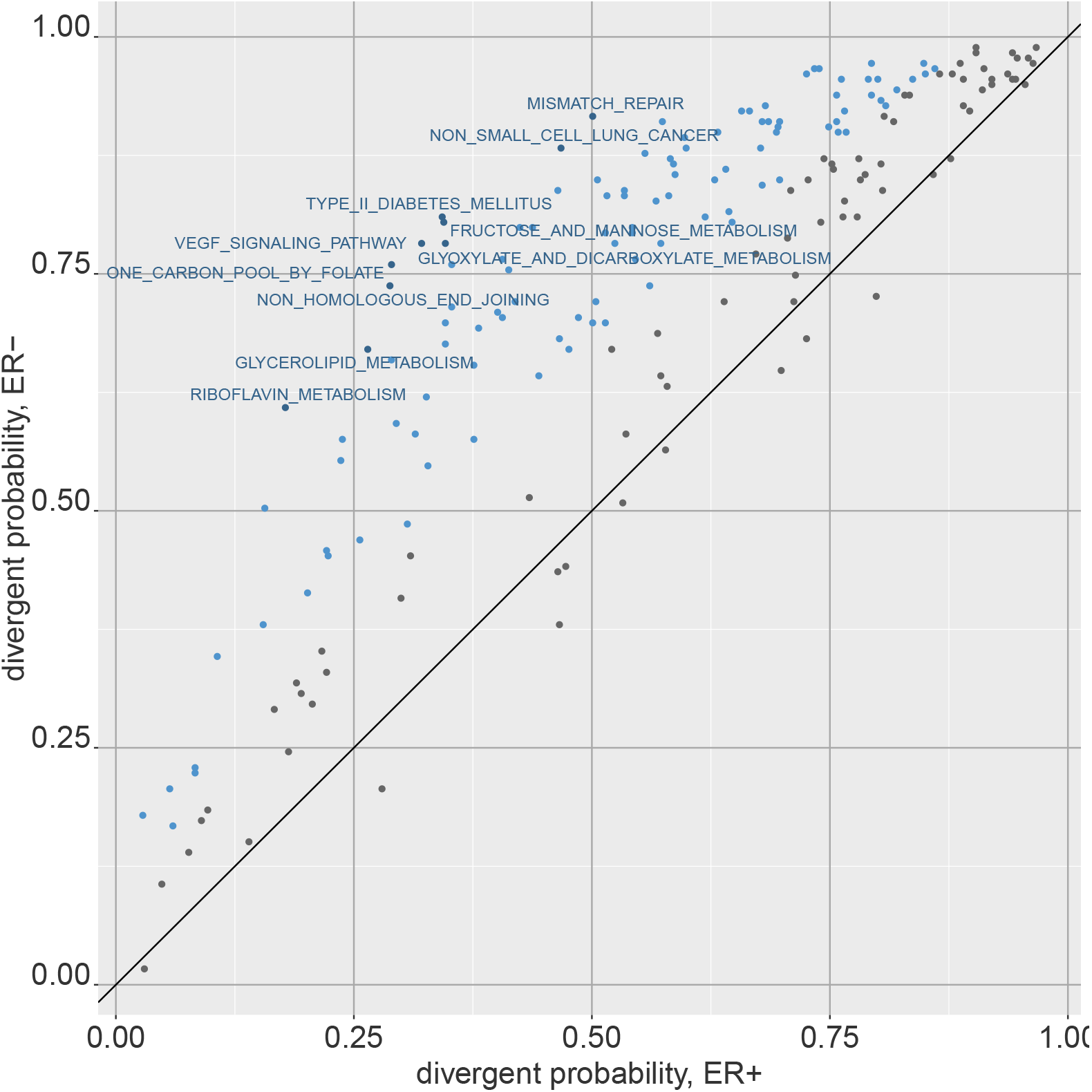
Differentially divergent gene sets between phenotypes. Similar to the univariate scenario, we can compare the divergence probability of multivariate features between groups of samples. The divergent probabilites among the ER+ and ER− breast tumor samples are shown here for KEGG gene sets, which the points in blue indicating meeting a bonferroni adjusted *P* ≤ 0.05 threshold from a *χ*^2^ test.

### Further Examples

In this section we provide a series of different example analyses that use RNA-Seq gene expression, microarray gene expression, and methylation 450k data with both univariate and multivariate modes of divergence.

### Discrimination between Luminal sub types with KEGG pathways

In this example, we demonstrate using multivariate divergence coding to train a classifier that provides a mechanistically useful interpretation, and examine its robustness across other datasets. Continuing the example provided in the previous section, we use the multivariate divergence coding obtained for the breast cancer tumor samples with respect to a normal breast baseline, with KEGG pathway gene sets as our multivariate feature sets of interest.

Figure 9 shows principal component analysis (PCA) applied to the KEGG pathway divergence coding. We simply take the binary matrix of breast tumor samples with the divergence coding for each KEGG pathway and apply PCA analysis as we would with any other data matrix.

**Fig 9.**
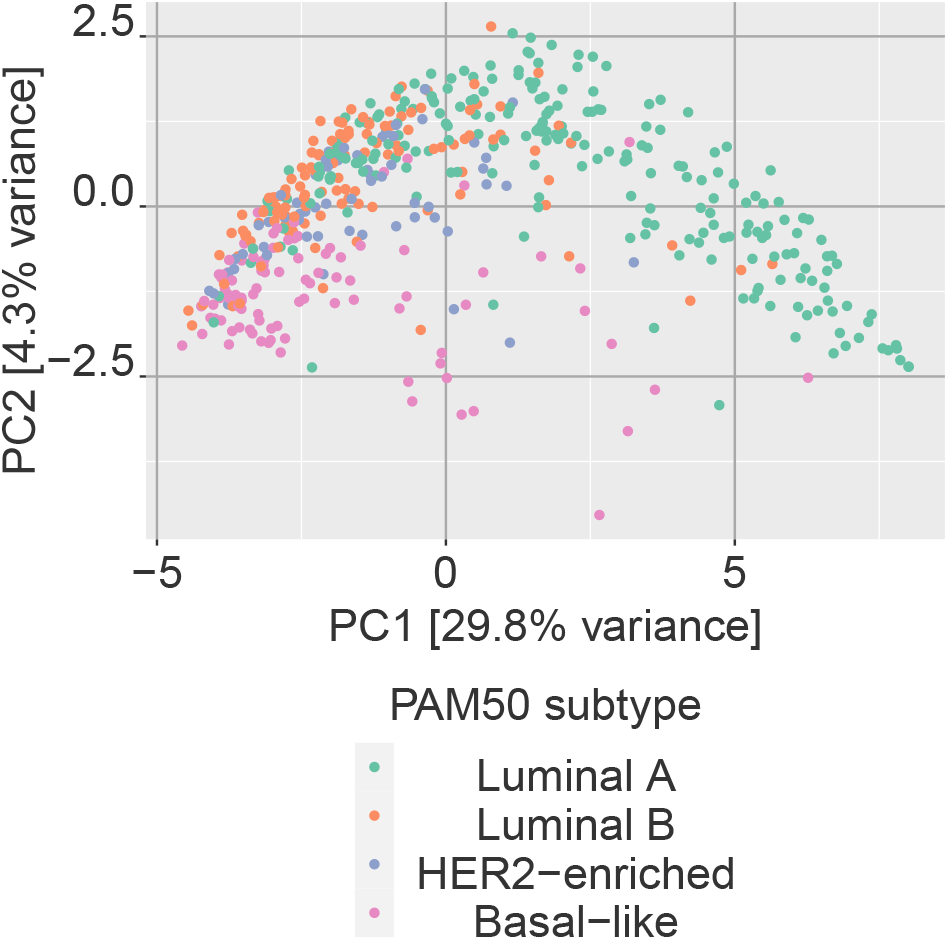
Principal component analysis of multivariate divergence. Digitized data can be analyzed with regular statistical tools such a principal component analysis (PCA). Here the first two PCs are shown for multivariate divergence values of the KEGG geneset collection for breast tumor samples with respect to a normal baseline.

The figure indicates that projection onto one of the leading principal components of the divergence data showed a significant ability to differentiate the Luminal A subset of tumor samples from the other PAM50 sub types showed. Based on this observation, we may train a classifier using the binary divergence coding for classifying between Luminal A and Luminal B sub types. Here we opt for decision trees and random forests as our classifiers of interest, given that with binary divergence data, these classifiers can provide more intuitive understanding of the biological underpinning behind the separation of the two groups, especially decision trees. Each branch of a decision tree is a binary decision of whether a certain KEGG pathway is displaying aberrant activity or not as indicated by divergence.

Candidate features for the classifiers were selected as the top 20 pathways resulting from *χ*^2^-tests between the two groups in the TCGA data (231 Luminal A samples and 127 Luminal B samples). The resulting decision tree provides a training accuracy of 0.85, while a random forest classifier provides 0.94. This decision tree is shown in figure 10.

**Fig 10.**
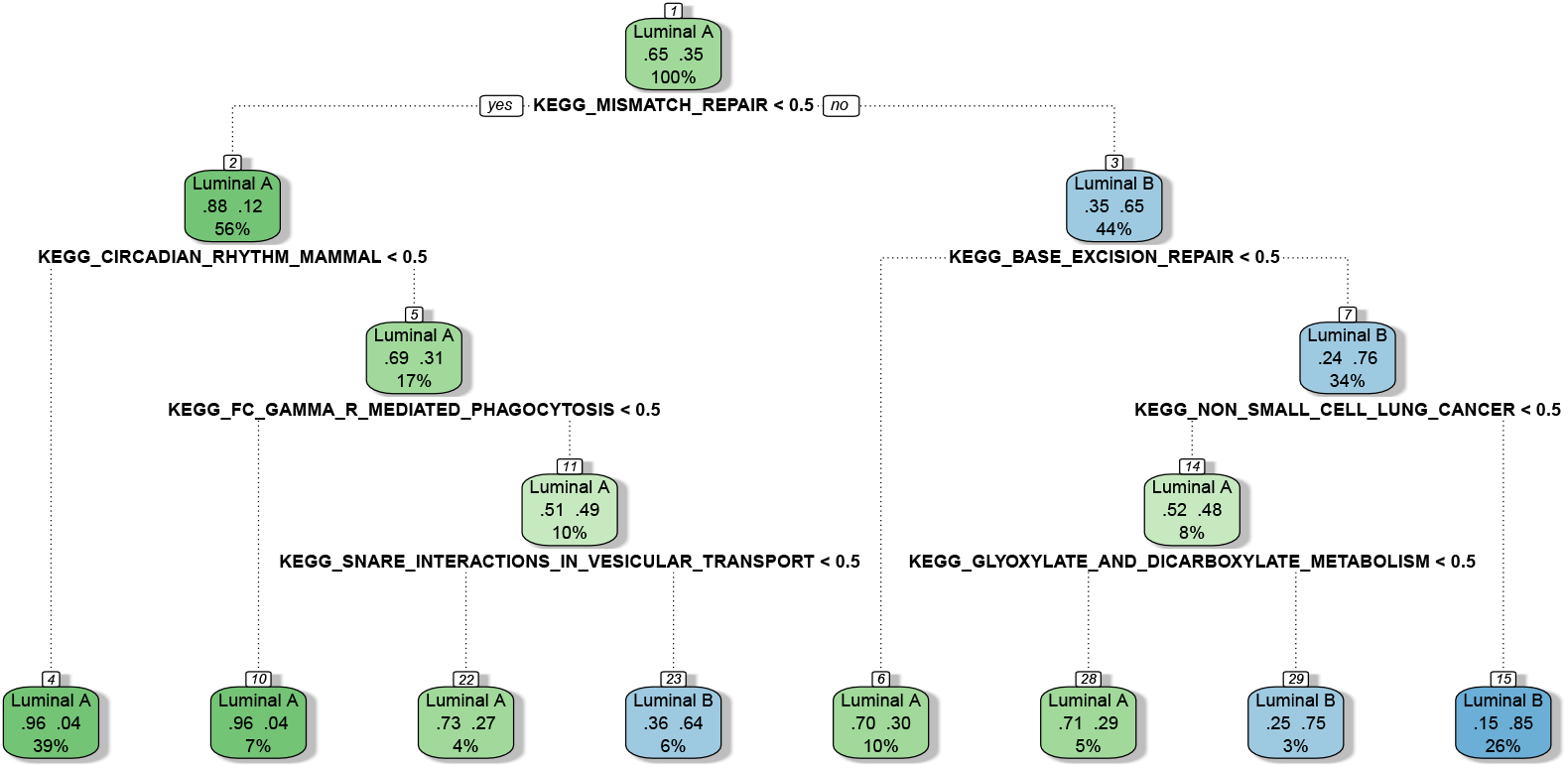
Decision tree for classification between Luminal A and Luminal B sub types. Using the divergence coding obtained for KEGG pathways, this decision tree was trained to separate Luminal A and Luminal B sub types. Each branch is a binary variable indicating whether a KEGG pathway is aberrant or not as indicated by multivariate divergence.

Next, we obtained independent microarray data for breast cancer from the GPL96 platform from publicly available studies (GEO accession numbers GSE2034, GSE12093, GSE17705, GSE25055, GSE25065) [10–13]. Due to lack of availability of normal samples from these studies, we used the ‘normal-like’ sub type as a baseline for obtaining KEGG pathway divergence coding for the Luminal samples. To ensure that the resulting data has a similar level of divergence to that from TCGA, we took the samples that would not be used for testing (i.e. omitting the Luminal samples), and computed divergence coding for the KEGG pathways for a range of *γ* values. From the resulting divergence codings, we selected the *γ* parameter that provided the highest correlation in divergence probabilities for the KEGG pathways to that from TCGA data, and used this setting to compute divergence codings for the Luminal A and B samples that would be used for testing (127 Luminal A samples, 66 Luminal B samples). Validating the above classifiers on these microarray samples provided accuracies of 0.64 and 0.71 from the decision tree and the random forest respectively (see Table 3).

**Table 3.** Classification results for Luminal A vs B with KEGG pathways. Classification results obtained for Luminal A and B samples from microarray data with the KEGG pathway based classifiers trained on TCGA data.

| classifier | data | accuracy | sensitivity | specificity | balanced accuracy |
| --- | --- | --- | --- | --- | --- |
| decision tree | train | 0.85 | 0.79 | 0.89 | 0.84 |
|  | test | 0.64 | 0.46 | 0.74 | 0.60 |
| random forest | train | 0.94 | 0.94 | 0.94 | 0.94 |
|  | test | 0.71 | 0.56 | 0.79 | 0.68 |

We believe with larger datasets and more complex training and tuning procedures, more robust classifiers that work well in cross-study cases may be developed. Our aim here is to demonstrate a fairly simple example of deriving a classifier from divergence that provides some understanding of the possible biological mechanisms underpinning the classifier, while also providing some degree of portability to a different dataset for validation.

### Gene set enrichment

Traditional types of analysis can also be performed with divergence coding if so desired. Gene set enrichment analysis (GSEA), which is the standard approach to examining whether a given ranking of genes in enriched in a collection of gene sets, is one such method we showcase here. For this example, we use the univariate gene expression divergence coding computed for breast tumor samples, along with the chemical and genetic perturbations collection of gene sets from the MSigDB project [9].

As a simple example, we use the ER+ and ER− sets of samples used previously. A noteworthy point here is than unlike a traditional setting where differential expression analysis performed on a binary phenotype comparison is used for obtaining the gene rankings, with divergence any set of features can be ranked over a single set of samples without the need for a binary comparison. Thus, we compute the divergence probabilities for genes for ER+ and ER− samples respectively, each set of samples providing us a ranking of genes. These rankings are then used to perform two sets of GSEA analyses, and the comparison of the results can be used to identify gene sets that are enriched with different directionalities in the two groups (see Figure 11). While in these results we compare the two ER groups, any ER group, or any other set of samples-may be analyzed with GSEA using divergence without the need for a binary phenotype comparison.

**Fig 11.**
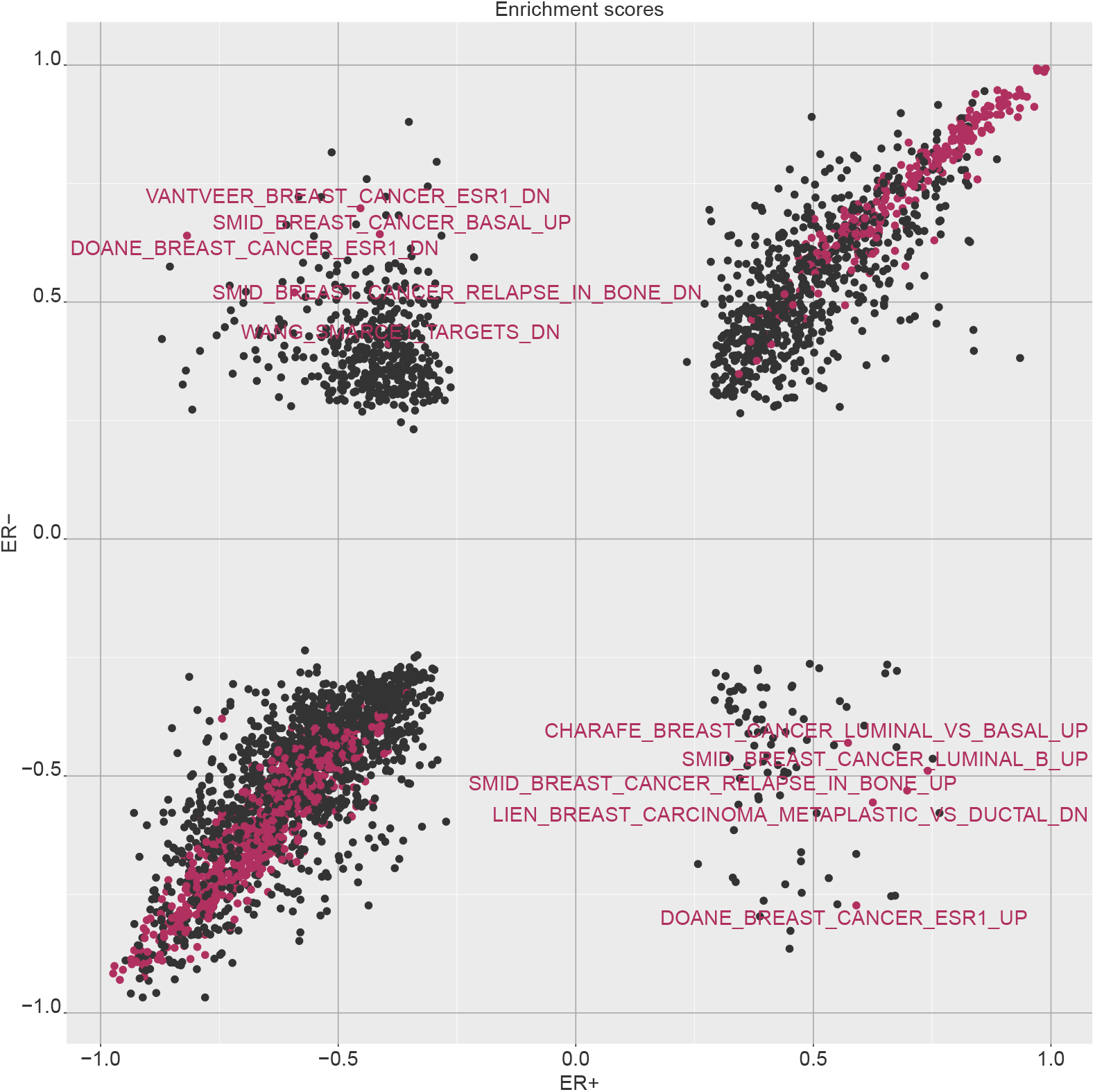
Comparison of GSEA results for ER+ and ER− samples with MSigDB chemical and genetic perturbation gene sets. The plot shows the enrichment scores obtained for the chemical and genetic perturbation gene sets by running gene set enrichment for ER+ and ER− groups separately with divergence coding. Gene sets of significant enrichment by FDR are colored, and those with opposing directions of enrichment in the two groups are labeled.

### Divergent CPG clusters

Here we use methylation data from the TCGA breast cancer cohort with divergence. The data we use are the methylation beta values in the [0,1] range indicating the proportion of methylation observed at a given CpG. As before, we use the methylation data for the TCGA normal samples as a baseline to compute divergence for the tumor samples. The mode of divergence used here is univariate divergence - i.e. we obtain a divergence coding for each sample at each CpG.

We can now use this data to look for differentially methylated regions. By arranging the CpG positions along the genome by chromosome and location, we may cluster the CpGs into regions of interest. Here we cluster CpGs in each chromosome into clusters such that that all adjacent CpGs in a cluster are no more than 300 bp apart. Then we simply compute the average sum of divergence observed over the tumor samples in each cluster. Suppose we have *k* clusters; given *i* = 1..*n* samples, for cluster *r_k_* this is

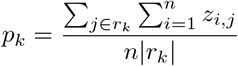

with *z_i,j_* being the binary divergence coding for CpG *j* in the cluster and sample *i*.

Further, we can assign a p-value to each region by computing a null distribution for the clusters through permutation of the CpG labels. The following CpG clusters appear to be the most highly divergent in breast tumor samples (Table 4).

**Table 4.** Highly divergent CpG clusters. The 10 CpG clusters with highest amount of divergence in the cluster.

| chromosome | start | end | cpgs | genes |
| --- | --- | --- | --- | --- |
| chr20 | 57425978 | 57427975 | 59 | GNAS-AS1,GNAS |
| chr6 | 33286243 | 33289721 | 55 | ZBTB22,DAXX,TAPBP |
| chr7 | 94285269 | 94287244 | 62 | SGCE,PEG10 |
| chr13 | 78491981 | 78494464 | 48 | RNF219-AS1,EDNRB |
| chr7 | 130129945 | 130132455 | 48 | MEST,MESTIT1,MIR335 |
| chr6 | 33130695 | 33134327 | 44 | COL11A2 |
| chr7 | 27183132 | 27184823 | 44 | HOXA-AS3,HOXA5 |
| chr4 | 154710223 | 154712582 | 36 | SFRP2 |
| chr11 | 14993377 | 14995756 | 39 | CALCA |
| chr1 | 92949336 | 92950838 | 31 | GFI1 |

We can compare these results to those obtained via bumphunter [14], a popular package used for finding differentially methylated regions. Figure 12 shows the cluster divergence sums (prior to normalization by cluster size) obtained for the top 10,000 clusters against the areas computed from bumphunter with the methylation data. Bumphunter searches for ‘differentially methylated regions’, or DMRs, which are regions within the specified clusters where all adjacent CpGs are differentially methylated between normal and tumor. For each cluster we take the DMRs found by bumphunter for that cluster and take the area (which indicates the degree of differential methylation as measured by a regression coefficient in the bumphunter model, summarized for the region) for the most significant DMR therein for comparison with our divergence based approach. The results show a strong agreement for the clusters with high amounts of divergence.

**Fig 12.**
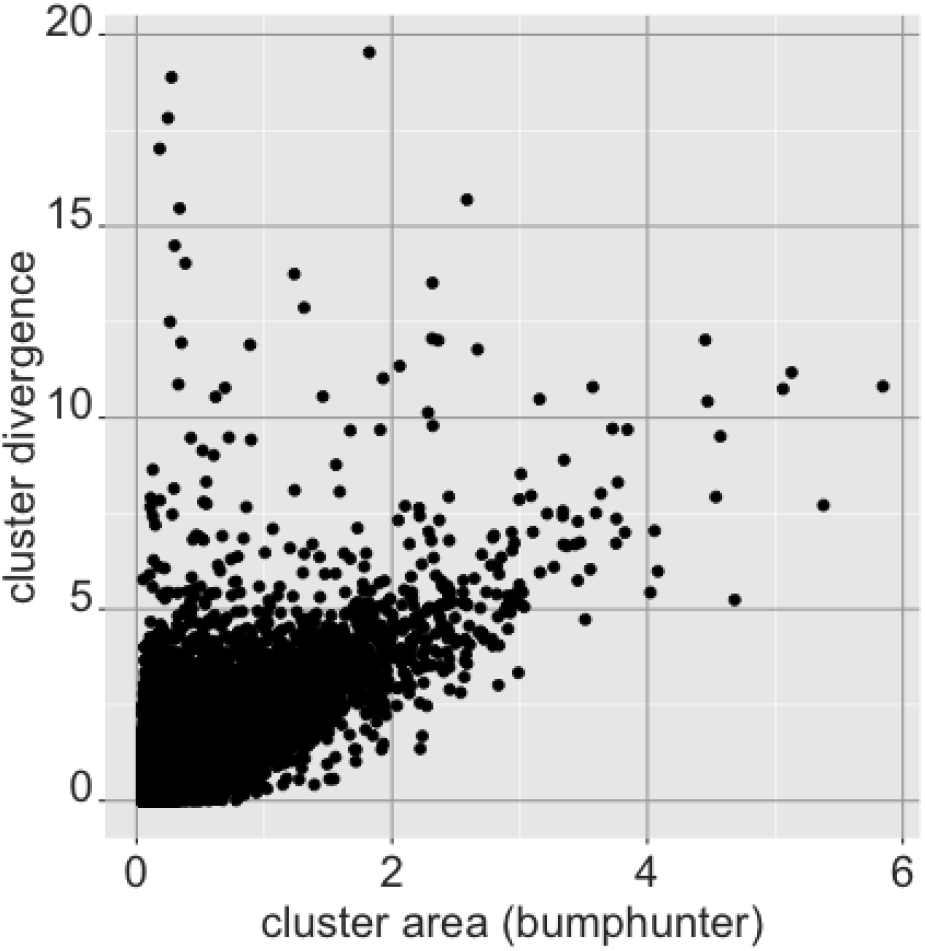
Comparison of cluster results from divergence against bumphunter for top CpG clusters. The y-axis shows the sum of divergent CpG instances in the cluster (standardized by the sample size). The x-axis show the area statistic obtained from bumphunter for the differentially methylated region mapped to that cluster, showing concordance between the two methods (correlation r=0.75).

### Significant gene-CpG associations

Having computed univariate divergence for two omics modalities - both methylation and gene expression - in this example we combine the two to discover what CpG - gene pairings show significant simultaneous divergence from both modalities. We are provided a table of CpGs mapped to nearby genes such that each CpG is mapped to a gene (with a single gene being mapped to one or more CpGs). We can compute the proportion of samples for which a given CpG - gene pair are both divergent.

For a given sample *i* (*i* = 1..*n*), suppose 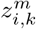 is the (binary) divergence value for CpG *k* and 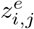 is the (binary) divergence value for gene *j*. Then the proportion of samples for which there is aberrant activity simultaneously in both modalities is an indication of the expression and the methylation between gene *j* and CpG *k* being related and co-aberrant. For gene *j* and CpG *k*, this is simply

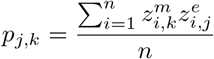

By considering the ternary form of divergence (i.e. −1/0/1), in addition to this proportion we can also compute the ‘concordant’ and ‘discordant’ proportions - i.e. the proportions of samples for which the sign of divergence are also in agreement. Thus samples for which both the gene and CpG are divergent with a +1, or both with a −1 - would be considered concordant, while other samples would be considered discordant.

Table 5 shows the 15 gene - CpG mappings with the largest measurements of co-divergence proportions. Further, we compared the values of *p_j,k_* with the correlation for the gene - CpG pair between the gene expression and methylation values. Figure 13 shows the results for the 10,000 gene - CpG pairs with the largest *p_jk_* values, indicating pairs with extremely positive or negative correlations are more likely to indicate high divergence. Some of these pairs are labeled with the associated gene. Figure 14 shows these pairs with the distance between the CpG and the associated gene as provided by the annotation, showing the CpGs mapped to a position near the gene are more likely to indicate co-aberrant activity as expected.

**Table 5.** Gene - CpG pairs showing high degree of simultaneous divergence. 15 Gene - CpG pairs with high proportions of samples being co-divergent are shown, with the chromosome and CpG locations. The concordant and discordant sample proportions (i.e. samples for which divergence in gene expression and methylation are either in the same direction or opposite directions) are also shown. Many of the high ranking pairs have discordant divergences.

| cpg | gene | proportion | concordant proportion | discordant proportion | chromosome | cpg location |
| --- | --- | --- | --- | --- | --- | --- |
| cg00185066 | SPRY2 | 0.835 | 0.003 | 0.833 | chr13 | 80910762 |
| cg22369786 | SPRY2 | 0.754 | 0.011 | 0.742 | chr13 | 80911625 |
| cg08648138 | TGFBR3 | 0.741 | 0.006 | 0.734 | chr1 | 92247731 |
| cg07581623 | DPYSL2 | 0.738 | 0.000 | 0.738 | chr8 | 26498829 |
| cg17080882 | TGFBR3 | 0.738 | 0.003 | 0.736 | chr1 | 92191624 |
| cg18081258 | NDRG2 | 0.731 | 0.001 | 0.729 | chr14 | 21494160 |
| cg04477962 | METTL7A | 0.728 | 0.003 | 0.725 | chr12 | 51317374 |
| cg05386769 | TNS1 | 0.724 | 0.000 | 0.724 | chr2 | 218681557 |
| cg13470462 | BOC | 0.722 | 0.003 | 0.719 | chr3 | 112965786 |
| cg14577406 | NFIB | 0.710 | 0.015 | 0.695 | chr9 | 14317153 |
| cg09284275 | MYH11 | 0.693 | 0.005 | 0.688 | chr16 | 15923486 |
| cg20902783 | NDRG2 | 0.688 | 0.003 | 0.686 | chr14 | 21494070 |
| cg15227610 | CRYAB | 0.688 | 0.003 | 0.686 | chr11 | 111782015 |
| cg00172849 | COL11A1 | 0.686 | 0.681 | 0.005 | chr1 | 103574531 |
| cg13474848 | NFIB | 0.686 | 0.004 | 0.682 | chr9 | 14350790 |

**Fig 13.**
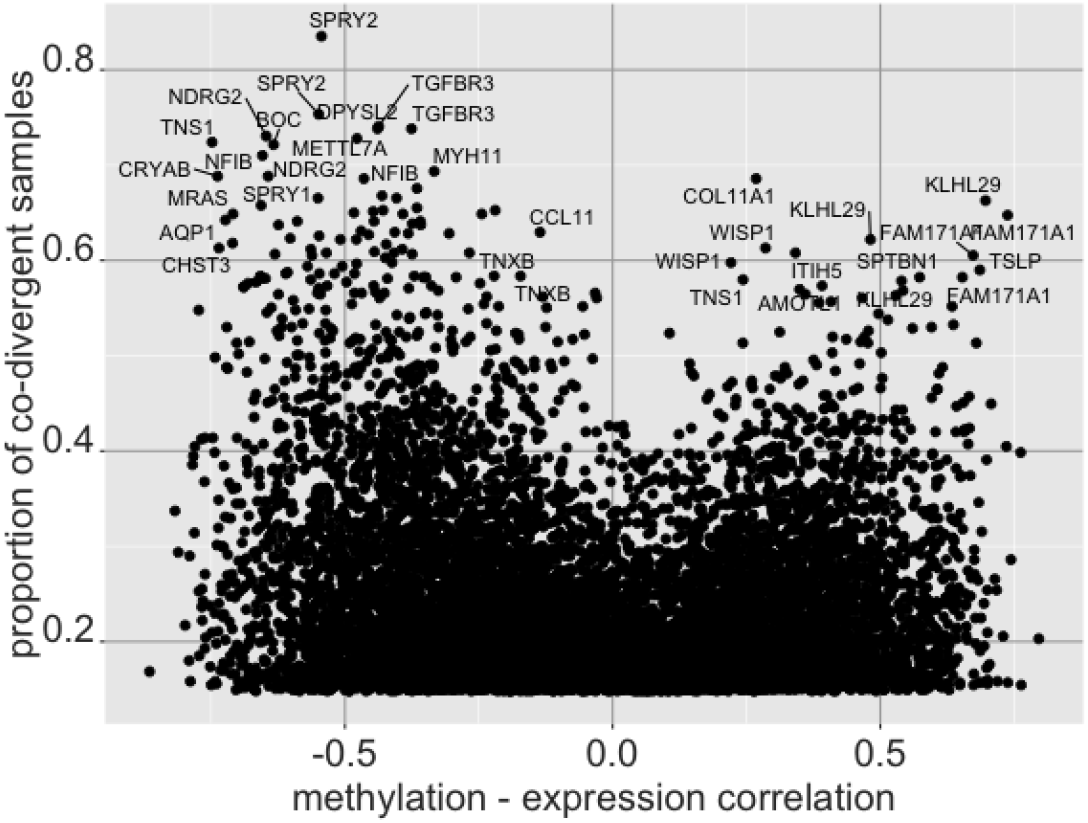
Comparison of gene - CpG pair co-divergence with expression-methylation correlations. The x-axis shows the correlation between the gene expression values and the methylations values for all samples, while the y-axis shows the proportion of samples for which both the gene and CpG are simultaneously divergent, with some of the high ranking pairs labeled by the participating gene.

**Fig 14.**
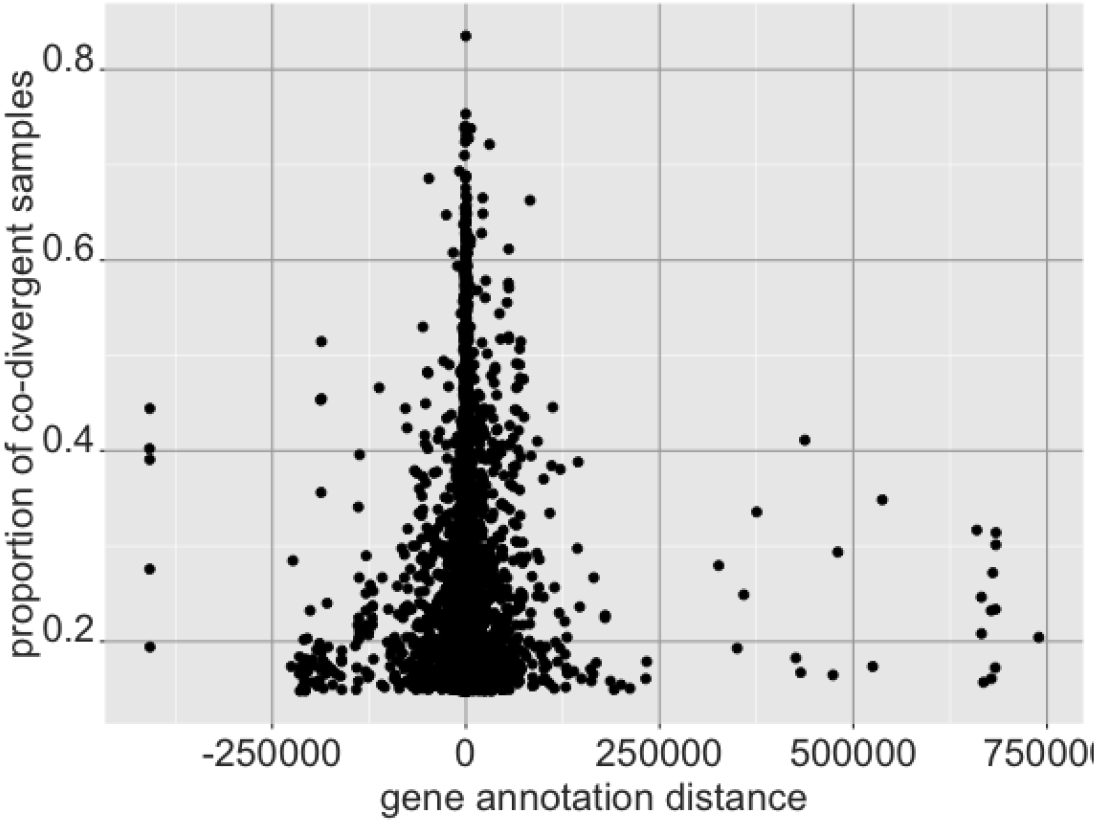
Comparison of gene - CpG pairs with gene annotation distance. The y-axis shows the proportion of co-divergence, while the x-axis indicates the distance from the CpG to the mapped gene. CpGs mapped closely to genes show high proportions of samples simultaneously divergent.

### Survival analysis with combined divergence

In this example, we compute and use co-divergence in the gene expression and methylation modalities in a different way. Using the gene - CpG annotation pairing previously mentioned, we use multivariate divergence to compute divergence for methylation data at the gene level. That is, sets of CpGs that are mapped to a single gene are treated as multivariate features of interest; multivariate divergence is computed for them using a normal methylation baseline. For genes mapped to a single CpG, binarized univariate divergence is computed. This provides us with a gene level divergence computed from the methylation data, which we can now combine with the gene level divergence computed previously with the gene expression data.

The two divergence codings for the same genes and samples - one from the methylation data and one from the gene expression data - can now be ‘added’. The binary values are combined to indicate co-divergence in both modalities. Thus for a given sample and gene, a 1 indicates the sample being divergent by gene expression level at that gene, and also being divergent in the CpG methylation space annotated to that gene (0 indicating no divergence in one or both modalities).

The co-divergence probabilities from this resulting data, for any gene considered, is an indicator of how likely the gene is demonstrating aberrant activity in both the transcriptome and the methylome simultaneously. The gene divergence probabilities resulting from this data are similar to the gene - CpG pair proportions computed in the previous example.

We now turn to look at relapse free survival (RFS) with this co-divergence data. We select tumor samples that are in stages I or II. This provides 23 samples with recorded relapses, and 260 censored samples. The selection criteria here is reflective of patients for whom existing genomic diagnostic tests such as MammaPrint [15, 16] are offered for risk evaluation.

To examine genes of interest with respect to relapse, we first select samples that have a censoring time of greater than 3 years among those with no relapse observed. Genes that have a divergence probability ≥ 0.3 in at least one group (i.e. between the relapsed or censored) are selected, followed by *χ*^2^-tests performed to select differentially divergent genes between the two groups. We select the resulting top 20 genes to fit to a cox proportional hazards survival model.

Genes are iteratively added to the model fit, keeping only genes that provide a significant coefficient estimate at a *P* ≤ 0.05 threshold. Table 6 shows the resulting estimates (log-rank test *P* < 10^-5^). Apolipoprotein D (APOD) is primarily down-divergent in the gene expression data, and low expression of it has been noted to be associated with poor prognosis in breast cancer [17,18]. Regulator of G protein signaling 6 (RGS6), which is down-divergent in most tumor samples, has been observed to be a suppressor of breast cancer [19]. BEX1 has been linked to ER+ breast cancer (a greater proportion of ER+ samples are divergent compared to ER− samples), as well as acute myeloid leukemia (AML), chronic myeloid leukemia (CML), and squamous cell cancer [20–22]. The human protein atlas data suggests PROZ as a prognostic marker in liver cancer, with higher protein expression observed in a subset of breast cancer samples [23, 24]. This suggests further examination of these genes both in the gene expression and methylation spheres for their contribution to increased risk in stage breast cancer.

**Table 6.** Cox proportional hazards model estimations with relapse free survival data. Genes were iteratively selected to add to a cox proportional hazards model with the combined gene expression and methylation divergence codings to obtain the above coefficients.

| gene | coef | hazard<br>ratio | se(coef) | z | p-value |
| --- | --- | --- | --- | --- | --- |
| APOD | 1.611 | 5.008 | 0.5 | 3.22 | 0.001284 |
| RGS6 | 1.037 | 2.82 | 0.478 | 2.17 | 0.030033 |
| PROZ | 0.972 | 2.642 | 0.453 | 2.146 | 0.031849 |
| BEX1 | 0.875 | 2.4 | 0.44 | 1.987 | 0.046873 |

## Conclusion

In the above results we have showcased a variety of analyses that can be performed with divergence at the univariate and multivariate level. While we have used RNA-seq, microarray and methylation 450k data here, the software is applicable to many other modalities of high dimensional omics data, some of which we have showed in [1]. While our previous publication was aimed at explaining the divergence framework from a statistical perspective, here we aim to explain how to use the package and conduct different types of analyses from divergence data with simple, practical examples.

Once the data has been processed as necessary and the baseline cohort identified, the R package can be used to compute the divergence coding quite easily. A full outline of the functions that can be used and the workflows possible are provided in the package vignette [4]. Here we have presented some of the many different ways that the digitized divergence coding can be visualized and analyzed by the user.

All the analyses conducted in this mansucript are available as R code through the publicly available github repository https://github.com/wikum/divergenceApplications. (The divergence package is available through https://www.bioconductor.org/packages/release/bioc/html/divergence.html.) The code provides an option for downloading the relevant TCGA data through the TCGAbiolinks package [25].

Divergence provides high utility with respect to omics data analysis. We note that it is a novel framework without any comparable prior methods: the ability to binarize omics data, applicability across many different platforms, enabling sample level scoring and analysis, and being applicable in both univariate and multivariate modes are highly desirable features that are not usually found together in other methods for omics data analysis. Given this fact, we believe that the divergence package will be highly useful to the computational biology research community in their pursuits.

## Acknowledgements

This publication was made possible though support from the NIH-NCI grants P30CA006973 and R01CA200859, and the Department of Defense (DoD) office of the Congressionally Directed Medical Research Programs (CDMRP) award W81XWH-16-1-0739.

## Declarations

The authors declare that they have no competing interests.

